# Differential regulation of eye specification in *Drosophila* by Polycomb Group (PcG) epigenetic repressors

**DOI:** 10.1101/2024.08.28.610158

**Authors:** Haley E. Brown, Brandon P. Weasner, Justin P. Kumar

## Abstract

During metazoan development, chromatin plasticity and genetic regulation are tightly connected to establish proper tissue fate and patterning. *Drosophila* imaginal discs are excellent models to study these processes as both genetic and mechanical injury can redirect their fate during regeneration. Reducing expression of *Polycomb* (*Pc*) results in the ectopic activation of the wing selector gene *vestigial* (*vg*) which in turn interacts with its DNA-binding partner Scalloped (Sd) – this forces the eye to transform into a wing. Reductions of other PcG members alone does not phenocopy this transformation. However, knocking down *Sex combs on midleg* (*Scm*) or *Scm-related gene containing four mbt domains* (*Sfmbt*) alongside the Pax6 gene *twin of eyeless* (*toy*) enables the eye-to-wing transformation. Using high throughput sequencing we show that *toy-Sfmbt* and *toy-Scm* knockdowns alter expression of wing selector and Hox genes, respectively. These findings provide new insights into how the fate of the eye is specified.

**Author Summary:** Distinct gene sets are differentially expressed in response to the knockdown of Polycomb Group (PcG) members suggesting multiple avenues may exist for the eye to be transformed into a wing.

## Introduction

A critical decision during metazoan development is the acquisition of tissue identity. Failure to correctly specify tissue fate can lead to the derailment of development and the onset of congenital disorders. As such, elucidating the molecular and developmental mechanisms underlying organogenesis is essential for understanding development and disease. The imaginal discs of the fruit fly, *Drosophila melanogaster*, are popular model systems for studying tissue growth, specification, pattern formation, and morphogenesis. These epithelial sacs originate during embryogenesis (Auerbach, 1936; Kaliss, 1939) and give rise to most of the external structures of the adult fly including the head, thorax, legs, wings, halteres, and genitalia (Geigy, 1931; Howland and Child, 1935). Because both genetic and mechanical injuries can redirect the fate of the imaginal discs (Weasner and Kumar, 2022), these tissues have become excellent model systems for identifying signal transduction pathways, transcription factors, and epigenetic regulators that guide developing tissues through a series of decision points as they make their way to adopting their final fate.

Studies in which cells from developing tissues were disassociated, transplanted into foreign hosts, and allowed to develop indicate that imaginal disc fate is determined during embryogenesis (Chan and Gehring, 1971; Gehring, 1970; Hadorn, 1968; Hadorn and Muller, 1974; Schubiger et al., 1969). Similar timings are also associated with the determination of tissue identity in vertebrates (Holtfreter, 1943; Moscona, 1957; Moscona and Moscona, 1952; Trinkaus and Gross, 1961; Weiss and Taylor, 1960). Over the course of three larval instar stages, the imaginal discs grow in size, are regionally specified, and undergo pattern formation. While many loss-of-function mutants result in the loss, reduced size, or structural distortion of imaginal disc derivatives, an array of genetic manipulations and mechanical injuries can also force an imaginal disc to undergo transdetermination, the process by which one disc loses its preprogrammed fate and adopts that of another. Each disc has a unique capacity to undergo transdetermination (Hadorn, 1968, 1978; Weasner and Kumar, 2022).

The phenomenon of transdetermination was discovered during studies aimed at understanding how and when the fates of imaginal discs are determined. In short, when whole imaginal discs are transplanted into larval hosts, they will autonomously develop, undergo metamorphosis, and produce adult derivatives that can be surgically isolated from the resulting adult hosts (Beadle and Ephrussi, 1935). The resulting adult derivatives are appropriately matched to the transplanted disc (i.e., a wing disc will produce an adult wing and notum). A similar phenomenon is seen when fragments of imaginal discs are transplanted (Hadorn, 1966, 1967). However, in a small number of instances the transplanted fragment will produce an inappropriate adult derivative. The term transdetermination was coined to describe these changes in fate (Hadorn, 1968). Some discs such as the antenna can take on the identity of multiple other discs while others like the eye disc having a much more limited capacity to have its fate altered (Hadorn, 1978; Weasner and Kumar, 2022). Here we focus on the development of the eye field and its sole propensity to transdetermine into a wing.

An understanding of the molecular mechanisms by which the eye primordium rejects the choice to develop into a wing during normal development is beginning to emerge. This is due in large part to a series of genetic manipulations that mimic the disc fragmentation studies and result in the eye-to-wing transformation. These include (1) double mutants of *eyes reduced* (*eyr*) and *eyeless* (*ey*) (Edwards and Gardner, 1966), (2) double mutants of *opthalmoptera* (*opht*) and *ey* (Goldschmidt and Lederman-Klein, 1958; Postlethwait, 1974), (3) double mutants of *lobiod* (*ld*) and *opht* (Kobel, 1968; Ouweneel, 1969, 1970), (4) simultaneous expression of the Hox gene *Antennapedia* (*Antp*) and activation of the *Notch* (N) pathway (Kurata et al., 2000), (5) combining the loss of *twin of eyeless* (*toy*) with ectopic *Antp* expression (Gehring et al., 2009), (6) forced expression of the histone modifying factor *winged eye* (*wge*) (Katsuyama et al., 2005; Masuko et al., 2018), (7) reduced expression of the *Polycomb* (*Pc*) epigenetic repressor (Brown et al., 2023; Zhu et al., 2018), and (8) forced expression of *vestigial* (*vg*) - a known wing selector gene (Kim et al., 1996; Simmonds et al., 1998). The roles that *Pc* and *vg* play in the eye-to-wing transformation are intertwined with each other as reductions in *Pc* expression lead to the upregulation of *vg* within the eye primordium (Brown et al., 2023; Zhu et al., 2018). This regulation is likely to be direct as two functional Polycomb Response Elements (PREs) are present within the *vg* locus (Ahmad and Spens, 2019; Herzog et al., 2014; Okulski et al., 2011; Ringrose et al., 2003). Our current model is that Pc epigenetically silences the *vg* locus within the eye during normal development thereby preventing it from adopting a wing fate (Brown et al., 2023). At the same time, the retinal determination (RD) network, led by the Pax6 transcription factors Ey and Twin of Eyeless (Toy), then pushes the eye primordium towards a retinal fate (Kumar, 2010). Together, these two mechanisms specify the fate of the compound eye.

In this manuscript we continue our investigation into the role that PcG proteins play in the eye vs wing decision by addressing two observations from an earlier study (Zhu et al., 2018). The first observation is that, unlike Pc, knocking down expression of other Polycomb Group (PcG) members individually does not induce an eye-to-wing transformation. This is unexpected as all PcG members are thought to work together to remodel chromatin and silence gene expression. Second, the eye-to-wing transformation can be induced if select PcG members are simultaneously knocked down with either *ey* or *toy*. This is reminiscent of the requirement for *ey* expression to be lost simultaneously with that of either *eyr* or *opht* in order for the fate of the eye to be redirected towards a wing (Edwards and Gardner, 1966; Goldschmidt and Lederman-Klein, 1958; Postlethwait, 1974). As both Ey and Toy are master regulators for eye specification and paralogs of each other (Czerny et al., 1999; Halder et al., 1995) our findings suggest that their loss enhances developmental plasticity of the disc which in turn enables transdetermination events.

Here we use RNA sequencing (RNA-seq) and <u>C</u>leavage <u>U</u>nder <u>T</u>argets and <u>R</u>elease <u>U</u>sing <u>N</u>uclease (CUT&RUN) to (1) understand why individual knockdowns of *Pc*, *Sfmbt*, and *Scm* have differential effects on eye fate, (2) investigate the impact that the loss of *toy* has on the transcriptome and chromatin, and (3) compare the molecular signatures of several genotypes that produce the eye-to-wing transformation (*Pc* knockdown, *toy-Sfmbt* double knockdown, and *toy-Scm* double knockdown). Our findings suggest that the eye-to-wing transformation in *Pc* and *toy-Sfmbt* knockdowns arises through a common mechanism; namely, the activation of the *vg* wing selector gene. In contrast, the *toy-Scm* double knockdown discs may use a different mechanism that is reliant on the activation of Hox genes. Our experimental data also suggest that Toy plays a role in maintaining a robust pattern of H3K27me3 at the *vg* locus during normal eye development. Levels of this repressive mark are dampened within the eye-antennal discs whenever *toy* is knocked down (either by itself or in combination with PcG members). Our findings shed new insights into the roles that Pax6 and PcG epigenetic regulators play in establishing the fate of the eye.

## Results

### Knocking down select PcG members alongside *toy* drives an eye-to-wing transformation

Our previous investigations into eye-antennal imaginal disc fate have shown that reducing *Pc* expression leads the eye to transform into a wing (Zhu et al., 2018) and in this context the reduction of *Pc* in the developing eye field relieves H3K27me3 repression of the *vestigial* (*vg*) locus (Brown et al., 2023). This allows for Vg to be expressed, co-opt its binding partner Scalloped (Sd), and force the eye to adopt a wing fate (Brown et al., 2023). Interestingly, although knocking down any of the remaining PcG members fails to drive this transformation the situation changes when these members are knocked down alongside *toy* (Zhu et al., 2018). For example, the combined loss of *toy* with *Scm-related gene containing four mbt domains* (*Sfmbt*) (Klymenko et al., 2006; Usui and Simpson, 2000) leads to a drastic eye-to-wing transformation that rivals that of the *Pc* transformation (Zhu et al., 2018). Here we ask if the eye-to-wing transformation occurs via the same downstream mechanism and investigate if and how *toy* mutants increase the developmental plasticity of the eye field.

To address these questions, we first combined the DE-GAL4 driver which is expressed in the dorsal portion of the eye (Morrison and Halder, 2010) with UAS based RNAi interference lines that target eighteen of twenty PcG members (Figure S1) and assayed the effects of losing individual PcG members on imaginal discs and adult heads. We looked for ectopic expression of the *Antp* Hox gene and/or a significant morphological change in the eye field as these are two hallmark features of *Pc* knockdown (PcKD) discs (Zhu et al., 2018). The DE-GAL4 driver itself has no effect on *Antp* expression nor on the morphology of the developing eye-antennal disc (Figure 1A). Ectopic *Antp* expression is detected when we knocked down five other PcG members: *pleiohomeotic* (*pho*), *Calypso*, *Sex combs extra* (*Sce*), *Sfmbt*, and *Sex comb on midleg* (*Scm*) even though the eye does not transform into a wing in these instances (Figure 1B-F). This confirms our earlier conclusion that while Antp may play a role in the eye-to-wing transformation, it does not drive this change in fate (Brown et al., 2023). phoKD, CalypsoKD and SfmbtKD discs appear morphologically wild-type and ectopic *Antp* expression is restricted to the peripodial epithelium of these discs (Figure 1B,C,E). The adult heads of phoKD and SfmbtKD animals also appear wild-type and even though CalypsoKD animals have lost some head capsule, their eyes, ocelli, and antennae appear normal (Table S1). SceKD discs activate *Antp* expression specifically within the antennal field and this results in its failure to be patterned properly (Figure 1D, asterisk, Table S1). And finally, ScmKD discs activate *Antp* in both the peripodial epithelium and disc proper of the entire eye-antennal disc. This leads to a slight increase in growth of the dorsal portion of the eye field (Figure 1F).

**Figure 1.**
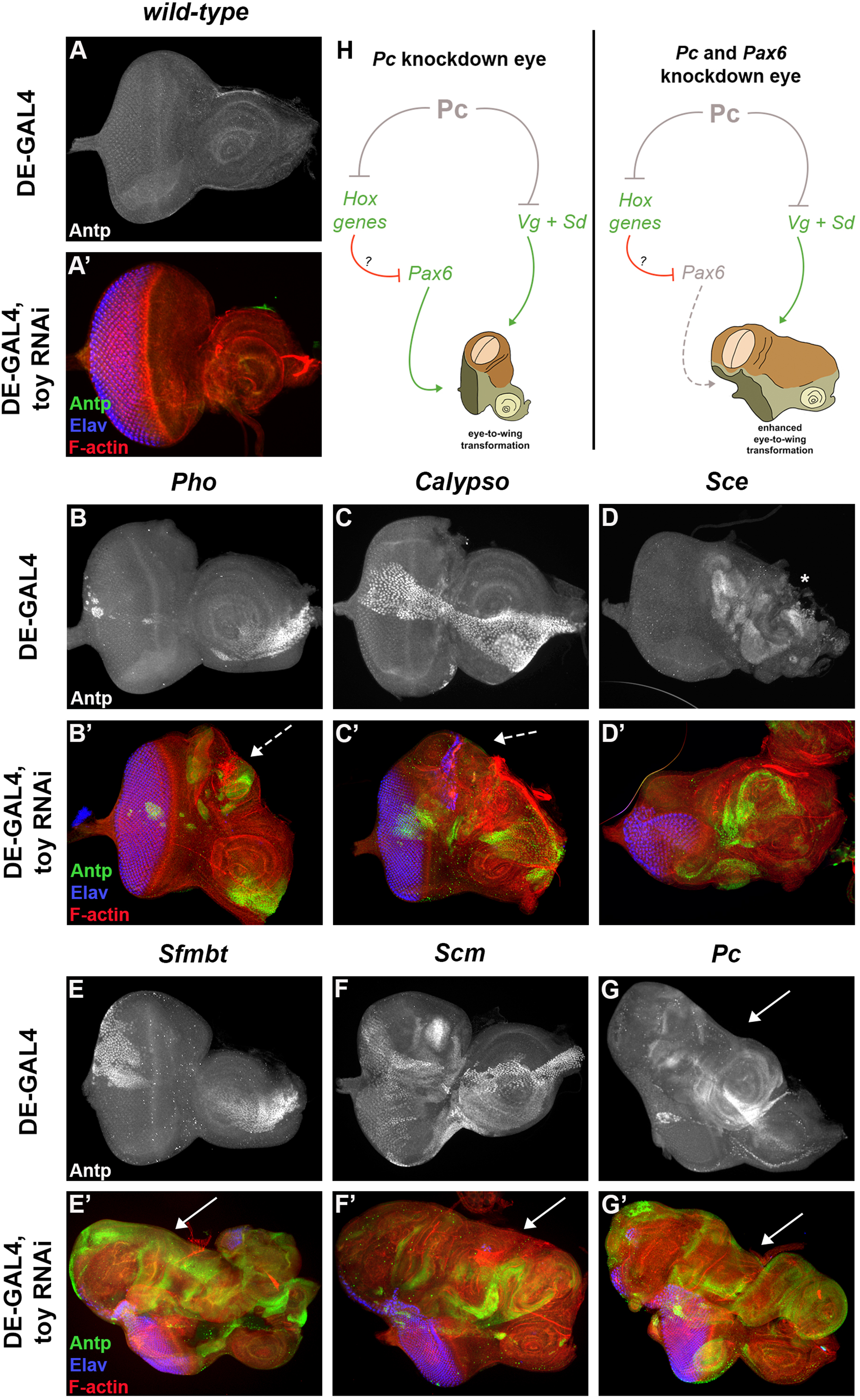
The knockdown of *Sfmbt* or *Scm* in addition to *toy* transforms the EAD. (A-G) Third-instar eye-antennal imaginal discs knocking down a PcG member with DE-GAL4; stained with Antp. (A’-G’) Third-instar eye-antennal imaginal discs knocking down a PcG member and toy with DE-GAL4; stained with anti-Antp (green), anti-Elav (blue), and phalloidin (red). Wild-type – DE-GAL4 (A) or DE-GAL4, toy RNAi (A’) – eye-antennal disc from a third-instar wandering larvae. Discs lacking either *Pho* (B), *Calypso* (C), *Sfmbt* (E), or *Scm* (F) alone ectopically express Antp, but otherwise maintain mostly wild-type morphology. The knockdown of *Sce* (D) reduces the eye and disrupts the antennal field (asterisk). Knocking down *Pc* with DE-GAL4 (G, arrow) grants an eye-to-wing transformation similar to that seen with *ey*-GAL4 (Brown et al. 2023). When combined with *toy* RNAi, knocking down *Pho* (B’) and *Calypso* (C’) leads to an outgrowth in the dorsal field (hatched arrows), while *Sce* (D’) shows similar eye filed disruption to DE>*Sce RNAi* alone (D). Knocking down *toy* and *Sfmbt* (E’) or *Scm* (F’) gives an exaggerated eye-to-wing transformation (arrows) like that seen in *toy-Pc* knockdown discs (G’). (H) Simplified model of the *Pc* eye-to-wing transformation (left) proposed in Brown et al. 2023, where the presence of Vg and Sd drives the wing transformation (brown) in the dorsal eye field, while continued presence of Pax6 maintains some eye fate (green). The combined loss of *toy* and *Sfmbt*, *Scm*, or *Pc* (right) leads to an enhanced eye-to-wing transformation more drastic than that of *ey>Pc RNAi* or DE>*Pc RNAi* alone. For all phenotypic scoring, n=30. Penetrance and severity of each phenotype are listed in Table S1.

Knocking down *Scm* is lethal at the late pupal phase and pharate animals most often fail to properly evolute their heads. Those that do show a potential wing-like outgrowth from the dorsal eye (Table S1). Overall these results show that the knockdown of *Pc* remains the only instance in which the loss of a PcG member, on its own, reproducibly drives the eye to transform into a wing (Figure 1G, arrow).

Since knocking down *toy* in conjunction with *Sfmbt* (toy-SfmbtKD) pushes the eye to adopt a wing fate (Zhu et al., 2018), we examined toyKD discs to see if the fate of the eye is affected and note that reducing *toy* expression, on its own, has minimal, if any, effects on the developing eye field (Figure 1A’). However, there are significant developmental consequences to the eye-antennal disc when the above six PcG members (*Sfmbt*, *pho*, *Calypso*, *Sce*, *Scm*, *Pc*) are lost in conjunction with *toy* (Figure 1B’-G’). In each instance there appears to be qualitatively more ectopic Antp protein and increased disc growth. For toy-phoKD and toy-CalypsoKD discs, the increased growth is restricted primarily to the dorsal eye field, while the remainder of the disc appears relatively normal (Figure 1 B’,C’, hatched arrows). toy-SceKD discs contain a second fully duplicated antenna, while the eye field is diminished in size (Figure 1D’). And quite surprisingly, toy-SfmbtKD, toy-ScmKD, and toy-PcKD discs all show a massive eye-to-wing transformation where the transformed wing tissue is no longer constrained to the dorsal eye field, but instead spans the entire dorsal region of the eye-antennal disc (Figure 1E’-G’, arrow). Since we have previously shown that the eye-to-wing transformation in PcKD discs is caused by the ectopic activation of the *vg* wing selector gene, we wanted to determine if the other toy-PcGKD transformations also rely on ectopic *vg* activation or whether sensitized backgrounds with reduced Pax6 (*toy*) levels engage another molecular mechanism entirely (Figure 1H). To investigate this question we used bulk RNA-sequencing (RNA-seq) and CUT&RUN to compare *Sfmbt*, *Scm*, and *Pc* individual knockdown discs to toy-PcG double knockdown discs.

Since the DE-GAL4 driver we used to express RNAi lines is expressed just within the dorsal-anterior region of the mid third larval instar disc (Figure S2A), we were concerned that small but statistically significant changes in gene expression within these cells may be masked by the remaining tissue that does not express the driver. To overcome this potential obstacle we attempted to use the Isolation of <u>N</u>uclei <u>Ta</u>gged in specific <u>C</u>ell <u>T</u>ypes (INTACT) (Deal and Henikoff, 2010; Ma and Weake, 2014) system to specifically isolate nuclei from genetically appropriate cell population for both RNA-seq and CUT&RUN. The version of the technique that we used in our study utilizes a UAS-GFP-KASH (<u>K</u>larsicht/<u>A</u>nc-1/<u>S</u>yne <u>h</u>omology) (Starr, 2011) transgene to label the outer nuclear envelope in a driver-dependent context. This allows for nuclei to be isolated using a combination of GFP antibodies and bead-based purification technologies. Driving UAS-GFP-KASH in the eye-antennal and wing discs with DE-GAL4 yields the expected expression patterns when only *toy* is knocked down (Figure S2A-B). However, incorporating the additional knockdown of *Sfmbt*, which triggers the eye-to-wing transformation, severely disrupts the GFP-KASH spatial expression pattern (Figure S2C-D, arrows). In control wing discs, the DE-GAL4 driver activates reporter expression only within the hinge, rather than the pouch (Figure S2B). When the dorsal eye field of the toy-SfmbtKD disc begins to transform into a wing, the expression GFP-KASH within the dorsal-anterior quadrant of the eye is abrogated and is activated just within the hinge domain of the ectopic wing. This leaves the ectopic wing pouch without GFP-KASH expression. As such, we abandoned using the INTACT system and instead focused on finding the time point at which the eye-to-wing transformation in each genotype has reached its zenith in terms of size. At these time points, the largest double knockdown discs, which are likely to have the most complete eye-to-wing transformation, would be selected for RNAseq and CUT&RUN analysis.

To determine the point in development at which each disc is at the apex of its transformation, we examined control, single knockdown, and double knockdown discs at various stages of third larval instar development (Figure 2). Wild-type animals, when reared at 25°C, complete larval development and begin pupariation around 120hr after egg lay (AEL). This is also the case for ScmKD and SfmbtKD discs in which ectopic *Antp* expression has been activated but the eye-to-wing transformation is not induced (Figure 2B-C, orange box). On the other hand, the onset of pupariation is delayed by twenty-four hours in PcKD, toy-PcKD, toy-ScmKD, and toy-SfmbtKD discs where the eye has transformed into a wing (Figure 2A, A’-C’, purple box). While PcKD and toy-PcKD discs consistently reach their peak size by 144hr AEL, toy-SfmbtKD and toy-ScmKD discs are on average smaller (Figures 1E’,F’, 2B’,C’). In fact, the eye-to-wing transformations are complete by 144hr AEL in only a small fraction of these two types of discs. We attempted to isolate toy-SfmbtKD and toy-ScmKD discs 12hrs (156hr AEL) and 24hrs (168hr AEL) later, but the vast majority of both types of larvae pupate by 156hrs. Wild-type discs continue to grow and undergo patterning during the first 12hrs of pupal development (Wolff and Ready, 1993). We speculate that this may also be the case for toy-SfmbtKD and toy-ScmKD discs as well. In the end, we decided to isolate discs from DE-GAL4 (control), toyKD, SfmbtKD, and ScmKD larvae that are 120hrs old while isolating discs from PcKD, toy-PcKD, toy-SfmbtKD, and toy-ScmKD larvae that are 144hrs old.

**Figure 2.**
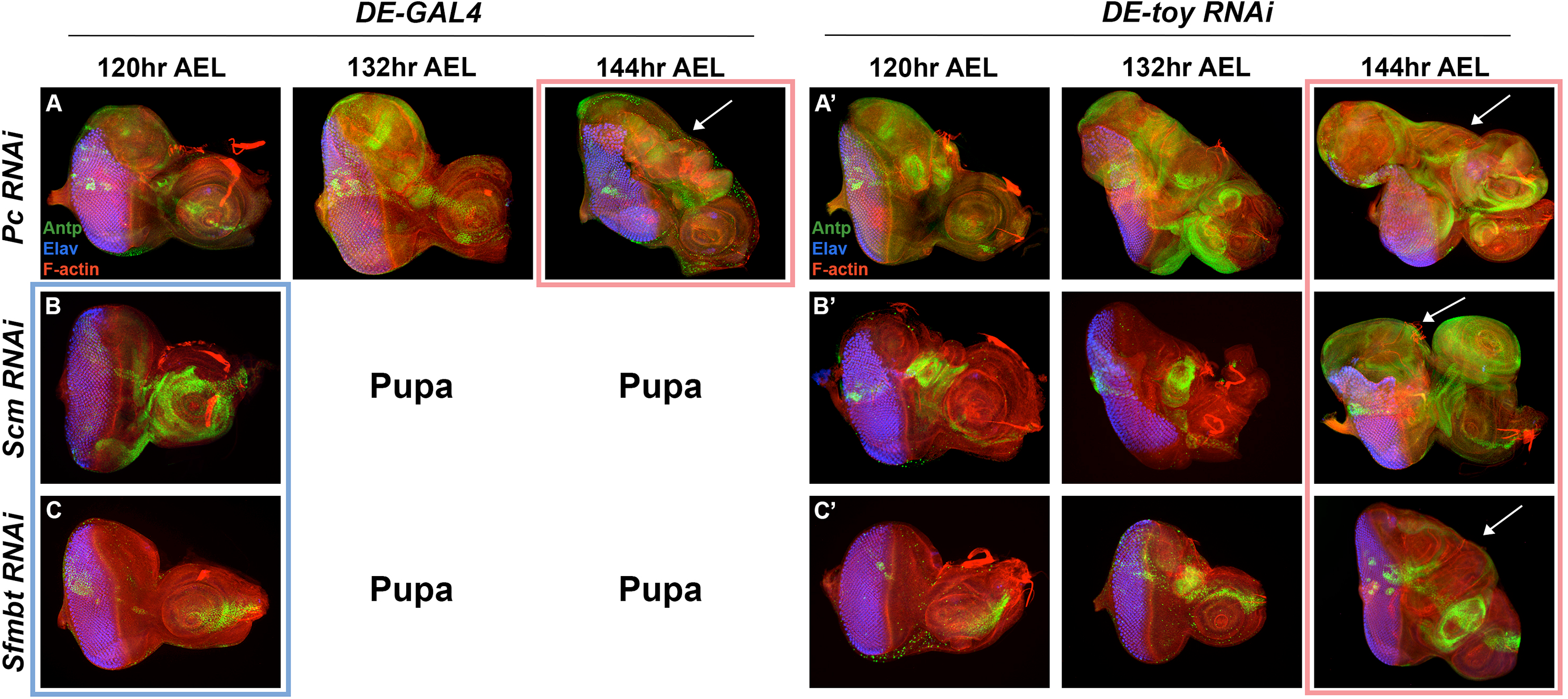
The eye-to-wing transformation occurs consistently at 144hr AEL. Time course of DE-GAL4 > *UAS-Pc Group* (*PcG*) *RNAi* (A-C) and DE-*toy RNAi* > *UAS-PcG RNAi* (A’-C’) at the end of third-instar development (120hr-144hr AEL); stained with anti-Antp (green), anti-Elva (blue), and phalloidin (red). When knocked down with DE-GAL4 alone, *Pc RNAi* animals delay pupation for an additional 24 hours and the dorsal eye field transforms by 144hr AEL (A). Development *Scm* (B) and *Sfmbt* (C) knockdown animals is not delayed, and these animals pupate after 120hrs with minimal eye-antennal disc transformation. Upon combined loss of *toy* with *Pc* (A’), *Scm* (B’), or *Sfmbt* (C’), all animals are developmentally delayed and transform by 144hr AEL. Arrows indicate an eye-to-wing transformation in the dorsal eye field. Boxed eye-antennal discs represent time points of transformed (purple) and non-transformed (orange) discs that were selected for RNA-sequencing.

### Bulk RNA-seq reveals few expression changes when *toy* alone is knocked down

To better understand the molecular mechanisms underlying the three robust instances of eye-to-wing transformation events, we first used bulk RNA-seq to compare the transcriptome profiles of PcKD discs to ScmKD or SfmbtKD discs that are morphologically wild-type and to toy-ScmKD or toy-SfmbtKD discs that harbor the eye-to-wing transformation (Table S2). Transcript counts show that *Pc*, *Sfmbt*, and *Scm* are reduced in the respective single and double knockdowns – this confirms that the RNAi lines that we are using do indeed knock down expression of their targets (Figure S3A-C). These counts also indicate that the knockdown of one PcG member does not affect the expression of other PcG members. This suggests that the phenotypes that we observe are due to the specific knockdown of *toy* and/or individual PcG members. Hierarchical clustering analysis shows that the transcriptomes of morphologically ‘wild-type’ eye-antennal discs (orange) discs tend to cluster together and are separated from the eye-antennal discs of those that have been transformed into a wing (purple) (Figure 3A). Interestingly, there are outliers with each of these two broad categories. One example is that two of three biological samples of SfmbtKD discs (which are morphologically wild-type) cluster with the genotypes that have the eye-to-wing transformation instead of the other genotypes that have morphologically normal discs. Another irregularity is that the three biological replicates of each genotype do not always cluster together (see replicates of the DE-GAL4 control discs). These minor anomalies may be due to the use of RNAi lines that can have a variable effect on transcript expression.

**Figure 3.**
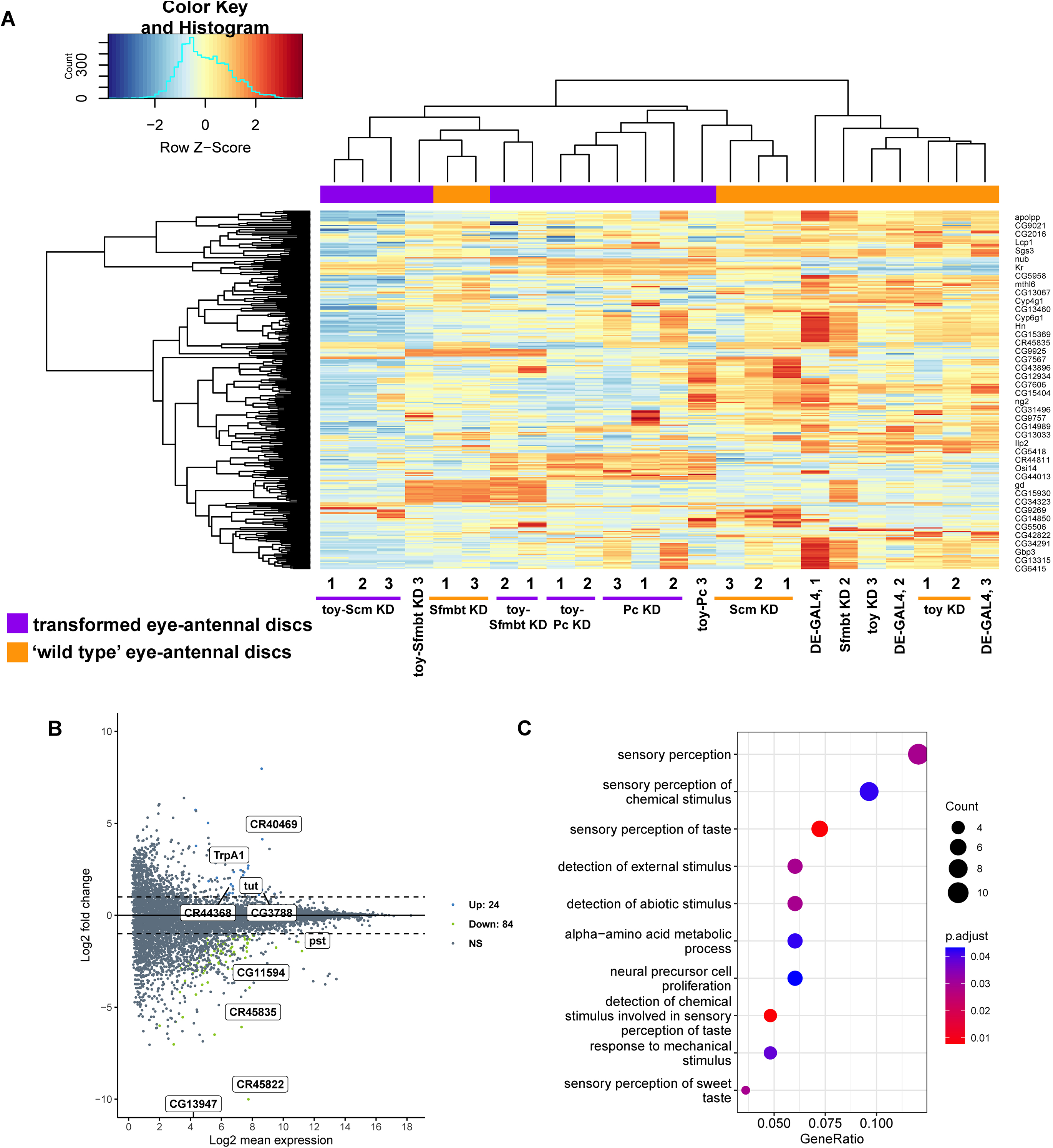
The eye-to-wing transformation clusters separately from morphologically wild-type eye-antennal discs. (A) CPM normalized transcript counts of 120hr AEL ‘wild-type’ eye-antennal discs (orange: DE-GAL4, DE-GAL4 > *UAS-toy RNAi*, DE-GAL4 > *UAS Sfmbt RNAi*, and DE-GAL4 > *UAS-Scm RNAi*) and transformed (purple: DE-GAL4 > *UAS-Pc RNAi*, DE-*toy* > *UAS-Pc RNAi*, DE*-toy* > *UAS-Sfmbt RNAi*, and DE-*toy* > *UAS-Scm RNAi*). Heatmap depicts hierarchical clustering of the top 100 most variable genes. (B) MA plot of differentially expressed genes between wild-type (DE-GAL4) eye-antennal discs and DE-*toy RNAi* (n = 108) eye-antennal discs. (C) Enriched Gene Ontology (GO) analysis of differentially expressed genes between wild-type (DE-GAL4) eye-antennal discs and ‘toyKD’ eye-antennal discs.

Since DE-GAL4, toyKD, ScmKD, and SfmbtKD discs are morphologically normal (Figures 1A,A’, 2B,C), it is not surprising that the transcriptomes of these control and mutant genotypes cluster together (Figure 3A). The relatively normal state of these discs is reflected in the small number and the nature of genes that are differentially expressed. This is best illustrated by the toyKD discs in which only 108 transcripts are differentially expressed when compared to DE-GAL4 control discs (Figure 3B). Most of these differentially expressed genes are unannotated protein-coding (CG) or non-protein-coding (CR) genes. The top Gene Ontology (GO) terms associated with these genes implicate them in the process of sensory perception processes (Figure 3C). As these genes are not expected to play roles during imaginal disc development it is no surprise that the discs are morphologically normal. Likewise, we do not observe drastic differences between the transcript profiles of PcKD and toy-PcKD discs (Figure S4A). Despite the increased level of tissue proliferation that occurs in toy-PcKD discs, only 69 additional genes are differentially expressed and most of these are annotated as being involved in cuticle development, sensory perception, or metabolic processes (Figure S4A).

This is not necessarily the case when comparing SfmbtKD or ScmKD non-transformed discs to their respective toy-SfmbtKD and toy-ScmKD counterparts, which is consistent with the eye-to-wing-transformation occurring within the double knockdown discs. When comparing SfmbtKD and toy-SfmbtKD discs to each other we note that an additional 209 transcripts are differentially expressed. While the top GO terms associated with these genes implicate them in small molecule biosynthetic and various metabolic processes (Figure S4B) we note that the wing selector gene *vg* is also upregulated in toy-SfmbtKD discs. Since *vg* is responsible for driving the eye-to-wing transformation in PcKD discs (Brown et al., 2023), our identification of *vg* here could be an indication that the eye-to-wing transformation in both PcKD and toy-Sfmbt knockdown discs both involve ectopic activation of *vg* and the formation of the Vg-Sd complex. In contrast, *vg* is not one of the 144 differentially expressed genes found in toy-ScmKD discs, which, are mostly annotated as being involved in protein folding (Figure S4B). Instead, several Hox genes including *Antp*, *Sex combs reduced* (*Scr*), *and Ultrabithorax* (*Ubx*) are differentially expressed. The differences between differentially expressed genes within toy-Sfmbt and toy-Scm discs may provide insight as to why the *toy-ScmKD* discs cluster separately from the toy-SfmbtKD and toy-PcKD discs (Figure 3A). It may also suggest that the eye-to-wing transformation in this context occurs via a mechanism that is independent of the Vg-Sd complex.

To better understand the potentially stepwise transcript changes underlying the eye-to-wing transformation we next made two additional transcriptome comparisons. The first was to compare the profile of DE-GAL4 control discs to those of single PcG knockdowns (Figure 4, Table S3). In our past study of PcKD discs we selected transcripts that were upregulated in both wild-type wing and PcKD transformed eye-antennal disc while being downregulated in the wild-type eye-antennal disc (Brown et al., 2023). Here we instead analyze all 645 genes that are differentially expressed in *PcKD* discs when compared to DE-GAL4 control discs (Figure 4A,B). In these discs several Hox genes including *Antp*, *Ubx*, *abdominal-A* (*abd-A*) and *Abdominal-B* (*Abd-B*) as well as numerous wing selection genes such as *vg*, *nubbin* (*nub*), *apterous* (*ap*), and *wingless* (*wg*) are upregulated while the Hox gene, *Deformed* (*Dfd*) is downregulated. Interestingly, despite the repression of eye development in the dorsal region of the disc, the expression of RD network members - including *toy* - is not significantly downregulated in these discs. This finding suggests that the loss or reduction of RD genes (and by extension eye fate) is not a requirement for the eye to transform into a wing. This is consistent with the ability of PcKD discs to undergo a fate transformation without the need for *toy* to be simultaneously knocked down.

**Figure 4.**
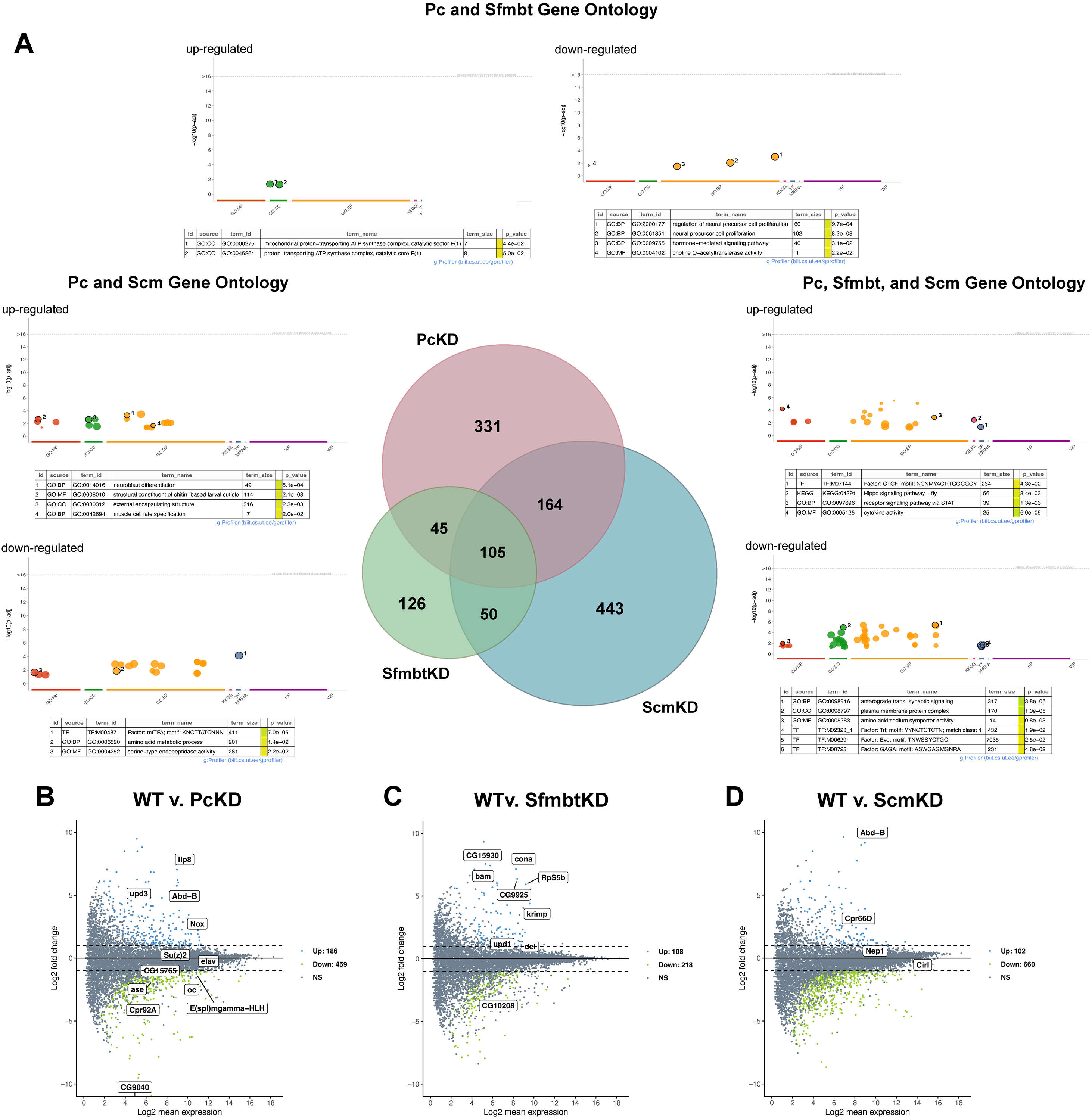
Transformed *Pc* knockdown discs share gene expression changes with *Sfmbt* and *Scm* knockdown discs. (A) Venn diagram showing overlap of differentially expressed genes (padj < 0.05) in *Pc* knockdown (PcKD, maroon, n = 645), *Sfmbt* knockdown (SfmbtKD, green, n = 326), and *Scm* knockdown (ScmKD, blue, n = 762) eye-antennal discs (middle). Up- and down-regulated genes in each category are listed in Table S3 (sheet 1). Gene Ontology (GO) analysis for genes shared between *Pc*, *Sfmbt*, and *Scm* (right), *Pc* and *Scm* (left), or *Pc* and *Sfmbt* (top). (B-D) MA plots of differentially expressed genes between wild-type (DE-GAL4) and *Pc* (B), *Sfmbt* (C), or *Scm* (D) knockdown eye-antennal discs – top differentially expressed genes are labeled.

Since the eye field does not transform in either SfmbtKD or ScmKD discs, we expected that fewer genes would be differentially expressed in these discs when compared to DE-GAL4 control discs. This is definitely true for SfmbtKD discs in which only 326 transcripts (vs 645 in PcKD discs) are differentially expressed between these two genotypes (Figure 4C). Interestingly, the minimal amount of ectopic *Antp* expression that we initially observed in these discs (Figure 1E) is not significant by DESeq2 parameters (padj < 0.05). Instead, the only Hox gene that is differentially expressed is *Abd-B*. In contrast, ScmKD discs unexpectedly displayed 762 differentially expressed transcripts when compared to DE-GAL4 control discs (Figure 4D). The loss of *Scm* within the eye field results in the mis-regulation of several Hox genes including *Dfd*, *Antp*, *Ubx*, *abd-A*, and *Abd-B* which is very similar to what we have observed for PcKD discs. Not surprisingly, neither *vg* nor any other wing selector gene are differentially expressed in either SfmbtKD or ScmKD discs. The lack of ectopic *vg* expression is the most likely explanation for why the eye fails to transform into a wing in these two PcG member knockdowns.

The second comparison that we made was between toyKD discs and toy-PcKD, toy-SfmbtKD, and toy-ScmKD discs (Figure 5, Table S3). Since the eye transforms into a wing in all three *toy-PcGKD* discs we expected that the three toy-PcG double knockdown genotypes would share a larger proportion of differentially expressed genes than the individual knockdown discs. However, this is not the case (compare Figures 4A to 5A, Tables S2,3) and we instead observe that slightly more differentially expressed transcripts are shared between all three individual PcG knockdown discs than the three toy-PcGKD discs. The transcript profiles of toy-PcKD discs differ from toyKD discs by the differential expression of 412 genes (Figure 5B). toy-PcKD show a similar differential regulation of Hox (up = *Scr*, *Antp*, *Ubx*, *Abd-B*; down = *lab*, *Dfd*) and wing (*vg*, *nub*, *ap*) genes. The transformed toy-SfmbtKD and toy-ScmKD discs also show drastically different expression patterns (314 and 800 genes respectively) when compared to that of the single toyKD disc. However, these two groups then seem to bifurcate, where toy-SfmbtKD discs mis-regulate the same set of wing genes as toy-PcKD discs, but only upregulate the *Antp* and *Abd-B* of Hox genes (Figure 5C). On the other hand, toy-ScmKD discs mis-regulate six of the eight Hox genes, but fail to differentially express *vg* (Figure 5D). These patterns of gene expression are interesting as they clearly suggest that there may be multiple molecular paths by which the eye-antennal disc can have its fate redirected towards a wing. This model is consistent with the relatively eclectic collection of genetic backgrounds (listed in the introduction) that produce an eye-to-wing fate change.

**Figure 5.**
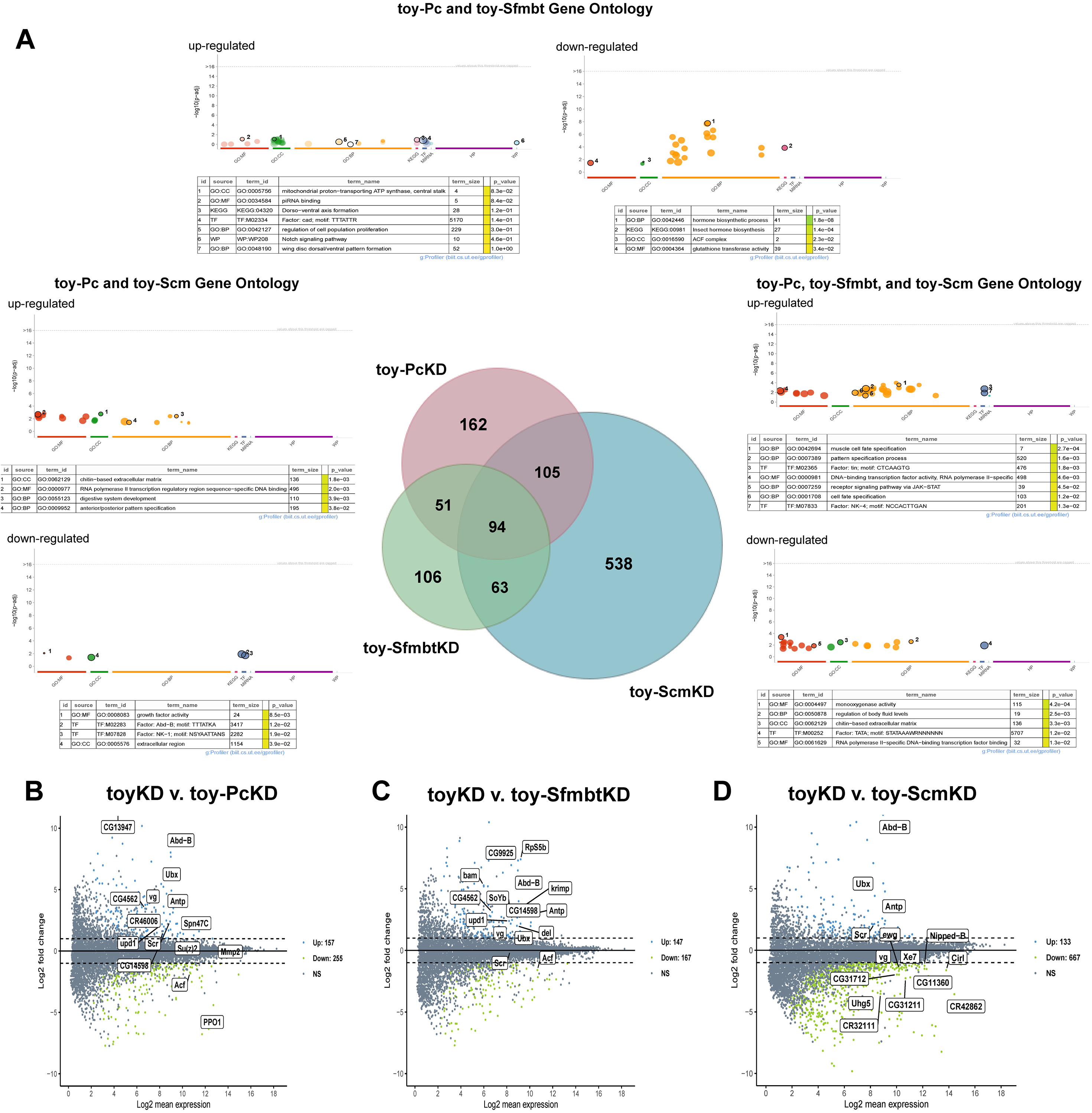
Transformed *Pc* knockdown discs are more similar to *Sfmbt* knockdown discs. (A) Venn diagram showing overlap of differentially expressed genes (padj < 0.05) in *toy-Pc* knockdown (toy-PcKD, maroon, n = 412), *toy-Sfmbt* knockdown (toy-SfmbtKD, green, n = 251), and *toy-Scm* knockdown (toy-ScmKD, blue, n = 800) eye-antennal discs (middle). Up- and down-regulated genes in each category are listed in Table S3 (sheet 2). Gene Ontology (GO) analysis for genes shared between *toy-Pc*, *toy-Sfmbt*, and *toy-Scm* (right), *toy-Pc* and *toy-Scm* (left), or *toy-Pc* and *toy-Sfmbt* (top). (B-D) MA plots of differentially expressed genes between *toy-Pc* (B), *toy-Sfmbt* (C), or *toy-Scm* (D) double knockdown discs alone and the toyKD discs. *vestigial* (*vg*) is differentially expressed in *toy-Pc* and *toy-Sfmbt* knockdown discs, but not *toy-Scm* knockdown discs.

### *PcKD and toy-SfmbtKD* discs share more similarities than *toy-ScmKD* discs

To determine if there is more than one path by which the eye can transdetermine into a wing, we compared the transcriptomes of PcKD discs to both *Sfmbt* and *Scm* single knockdown discs and toy-Sfmbt and toy-Scm double knockdown discs. Differential expression analysis (log2 fold change, adjusted p < 0.05) of these tissues identified 503 genes that were differentially expressed between PcKD and SfmbtKD, 458 genes between PcKD and ScmKD, 252 gene between PcKD and toy-SfmbtKD, and 657 genes between PcKD and toy-ScmKD (Figure 6A-F). This indicates that of all the tested comparisons, the transcriptomes of toy-SfmbtKD discs may be the most similar to that of PcKD discs, while toy-ScmKD discs may be the most distinct. This correlates with our observations that toy-ScmKD discs cluster separately from the other genotypes that produce an eye-to-wing transformations (Figure 3A).

**Figure 6.**
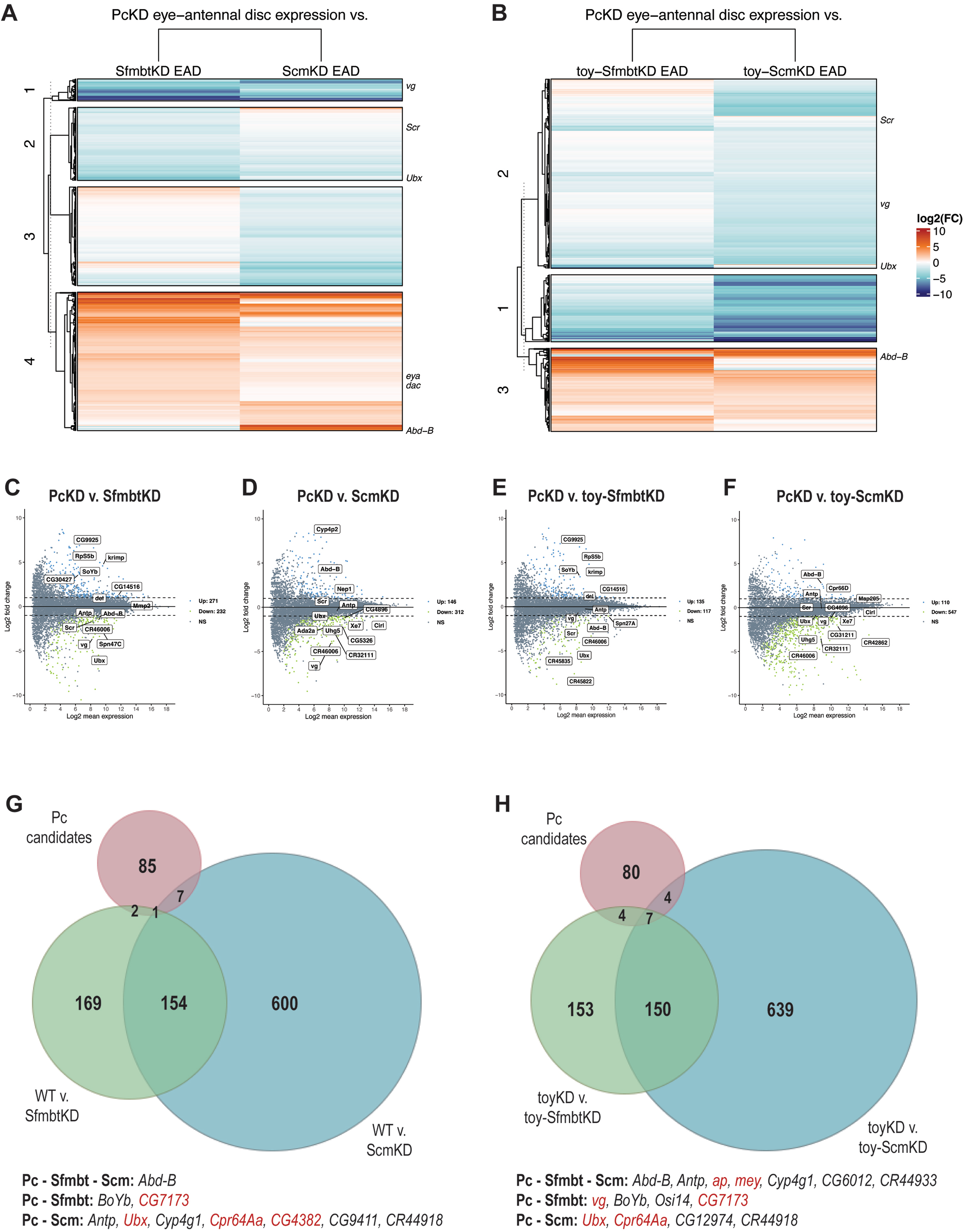
Previously identified eye-to-wing candidates are differentially expressed in *toy-Sfmbt* and *toy-Scm* knockdown discs. (A-B) Heatmap of log2 transformed differential transcript expression between *Pc* knockdown eye-antennal discs and either SfmbtKD and ScmKD eye-antennal discs (A) or toy-SfmbtKD and toy-ScmKD eye-antennal discs (B). Known eye genes, wing genes, and Hox genes are listed to the right. (C-F) MA plots of differentially expressed genes between *Pc* knockdown discs (PcKD) and SfmbtKD (C, n=503), ScmKD (D, n = 458), toy-SfmbtKD (E, n = 252), or toy-ScmKD (F, n = 657) discs. The top 10 differentially expressed genes as well as known wing, eye, and Hox genes are annotated. (G-H) Venn diagram of known *Pc* eye-to-wing candidates (maroon, Brown et al. 2023) compared to differentially expressed genes in SfmbtKD (green) and ScmKD (blue) discs (G) or toy-SfmbtKD (green) and toy-Scm (blue) discs (H). Overlapping candidates are listed below, with most likely candidates (shared between early, mid, and late development) highlighted in red.

In our previous study of the PcKD discs we identified 95 candidates that were differentially expressed during either early, mid, or late third-instar development (Brown et al., 2023). To determine if any of these candidates play roles in the *toy*-dependent eye-to-wing transformations, we asked if any of these genes are also differentially expressed in SfmbtKD and ScmKD discs (Figure 6G) or in toy-Sfmbt and toy-Scm discs (Figure 6H). Although 10 differentially expressed candidate genes are shared between either PcKD and either SfmbtKD or ScmKD discs, the only one that is differentially expressed in all three types of discs is *Abd-B*. Similarly, of the 15 differentially candidate genes that are shared between PcKD and either toy-SfmbtKD or toy-ScmKD discs, only the Hox genes *Antp* and *Abd-B* as well as the wing selector gene *apterous* (*ap*) are found in all three transformed discs (Figure 6H). These are unlikely to drive the eye-to-wing transformation as our previous work indicates that the forced expression of these Hox genes severely abrogates retinal development without inducting the eye to wing transformation. Likewise, ectopic expression of *ap* does not induce the eye-to-wing transformation either (Brown et al., 2023). The wing selector gene, *vg*, that we have shown to drive the *Pc*-dependent eye-to-wing transformation is only upregulated in toy-SfmbtKD discs (Figures 6H, red text, 7B). This may suggest that in both PcKD and toy-SfmbtKD discs the ectopic formation of the Vg-Sd complex drives the transformation of the eye into a wing while a distinct mechanism may drive this change in fate in toy-ScmKD discs. Of note, *Antp* expression is upregulated in toy-ScmKD discs. The combination of *Antp* expression and *toy* reduction is known to induce an eye-to-wing transformation (Gehring, 2009), thus it is also possible that this genetic combination is the underlying mechanism for the eye-to-wing transformation in toy-ScmKD discs. A second possibility is that the transformation of an eye into a wing may be driven by the combination of knocking down *toy* and upregulating *ap*. Ap sits upstream of *vg* within the wing gene regulatory network and Vg controls only a subset of all Ap functions (Delanoue et al., 2002). It is possible that the simultaneous loss of *toy* and upregulation of *ap* could bypass the requirement for the Vg-Sd complex and force the eye to transform into a wing.

### Reductions in *toy* dampen H3K37me3 marks at the *vg* locus

To gain insight into how Toy mediates the eye-to-wing transformation we used CUT&RUN to analyze H3K27me3 levels in eye-antennal discs of control and transformed larvae (Figure 7A,S5). We focused on the *vg* locus as it is activated in both PcKD and toy-SfmbtKD discs. H3K27me3 heavily decorates the *vg* locus in DE-GAL4 control discs (Figure 7A, first row). In PcKD discs this methylation mark is drastically reduced but not completely eliminated (Figure 7A, third row). As a result, *vg* expression is ectopically activated within the eye field (Figure 7B) (Brown et al., 2023; Zhu et al., 2018). Interestingly, H3K27me3 levels are significantly dampened when *toy* expression is knocked down alone (Figure 7A, second row). This results in a slight increase in *vg* expression (Figure 7B, left pair)) but no change in the fate of the eye (Figure 1A’) (Zhu et al., 2017). Moreover, H3K27me3 marks at the *vg* locus are depressed even further when toy-PcKD and PcKD discs are compared to each other (Figure 7A). This is consistent with an even higher increase in *vg* expression (Figure 7B, middle pair) and a more extensive eye-to-wing transformation in toy-PcKD discs (Figure 2A,A’). Finally, while the *vg* locus within SfmbtKD discs remains heavily decorated with H3K27me3 marks (Figure 7A, fifth row), those levels drop significantly in toy-Sfmbt KD discs (Figure 7A, sixth row). In fact, the levels of H3K27me3 in toy-SfmbtKD discs are even lower than those in toyKD discs (Figure 7A, second and sixth row). The further depression in H2K27me3 levels appears sufficient to allow for activation of *vg* expression in toy-Sfmbt discs (Figure 7B, right pair) and the eye-to-wing transformation (Figures 1E’,5C) (Zhu et al., 2018). As such, we conclude that a potential role for Toy (and possibly Ey) during normal eye development could be to participate in the maintenance of robust levels of H3K27me3 at the *vg* locus.

**Figure 7.**
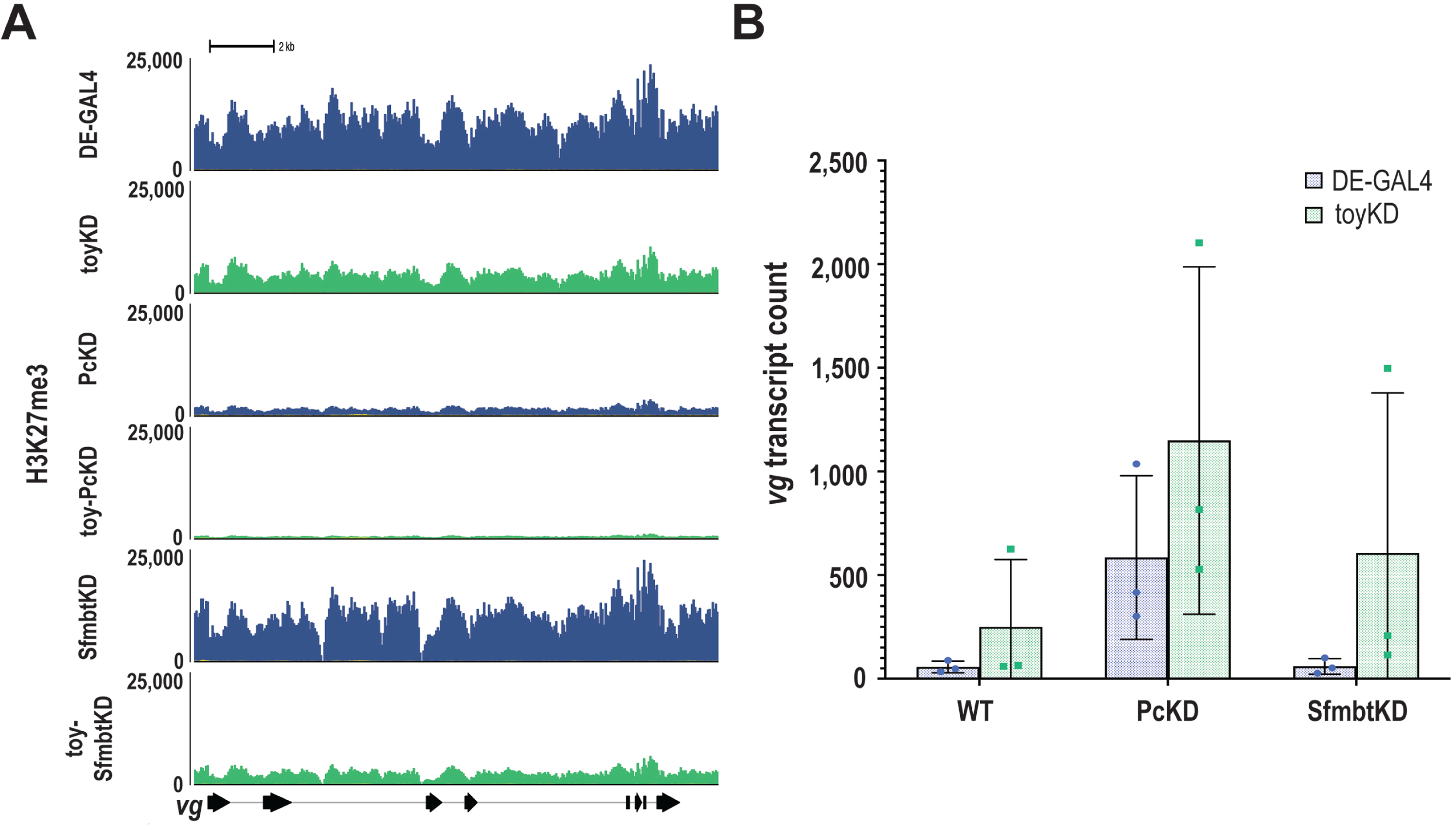
RNAi perturbation reduces H3K27me3 levels at *vg* locus. (A) CUT&RUN H3K27me3 peaks at the *vg* locus in either the ‘wild-type’ (DE-GAL4) eye-antennal disc (blue) or toyKD eye-antennal disc (green) crossed to *Pc* or *Sfmbt* RNAi lines. (B) Bar graph represents transcript counts of three replicates for the DE-GAL4, PcKD, or SfmbtKD (blue), and toyKD, toy-PcKD, or toy-SfmbtKD eye-antennal discs (green).

## Discussion

In this manuscript we have continued our investigation into the role that PcG epigenetic repressors play in regulating the choice between eye and wing fates in *Drosophila*. Our past studies have shown that in the absence of Pc expression within the eye field, H3K27me3 histone modifications across the *vg* locus are abrogated and *vg* is inappropriately expressed. This results in the ectopic formation of the Vg-Sd complex which then goes on to reprogram the eye into a wing (Brown et al., 2023; Zhu et al., 2018). In the course of these studies we made two unexpected observation. First, reducing the expression of other PcG members did not have the same effect on the fate of the eye. Second, the eye could be redirected towards a wing fate if the expression of select PcG members were simultaneously reduced along with that of the Pax6 genes *ey* and *toy*. Here we have focused on the mechanisms by which the eye transforms into a wing when *toy* levels are simultaneously reduced with either *Sfmbt* or *Scm* – two PcG members

We undertook this investigation because all PcG members are thought to function together to silence target genes (Figure S1). This is particularly true of *Pc*, *Sfmbt*, and *Scm*. *Sfmbt* and *pleiohomeotic* (*pho*) encode members of Pho-repressive complex (PhoRC) (Kassis et al., 2017). This complex binds to its molecular targets by recognizing PRE elements (Brown et al., 1998; Klymenko et al., 2006). PhoRC physically interacts with both PRC1, of which Pc is a member, and PRC2. PhoRC-PRC1 interactions are mediated by Scm which acts as physical bridge between Sfmbt and Polyhomeotic (Ph), a member of PRC1 (Frey et al., 2016; Grimm et al., 2009; Peterson et al., 1997). PhoRC-PRC2 interactions are mediated by physical contacts between Pho and Enhancer of zeste (E[z]) (Wang et al., 2004). Since PhoRC, PRC2, and PRC1 all converge at PRE sites, it is surprising that individual reductions in the expression of all three complex members do not have the same effect on the fate of the eye. Instead our observations suggest that eye specification is differentially regulated by individual PcG proteins. Our transcriptome analysis of *Pc*, *Sfmbt*, and *Scm* single knockdown discs confirms this hypothesis as its shows that expression of the *vg* wing selector gene is upregulated only in PcKD discs and not within either SfmbtKD or ScmKD discs.

We also attempted to address the role that Toy plays in maintaining the fate of the eye field. We first asked whether Toy plays a role in regulating *vg* expression either on its own or in combination with PcG members. Toy appears to function solely as a transcriptional activator (Weasner et al., 2009) which is inconsistent with a potential role in repressing *vg* expression in the eye disc. Indeed, our RNA-seq data indicates that *vg* expression is not differentially expressed in toyKD eye-antennal discs. While this data on its own would indicate that Toy does not regulate *vg* expression, we did find two instances in which the loss of Toy did paradoxically result in an upregulation of *vg* expression. First, while *vg* expression is not differentially expressed in SfmbtKD discs, it is ectopically expressed at significantly higher levels in toy-SfmbtKD discs. Second, although *vg* is differentially expressed in PcKD discs, its expression levels are even higher in toy-PcKD discs. How can the transcriptome results of toyKD discs be reconciled with those of toy-SfmbtKD and toy-PcKD discs? The answer may lie within our CUT&RUN data on H3K27me3 histone marks. As noted above, reducing *toy* expression in either single of double knockdown conditions results in a dampening of the H3K27me3 mark at the *vg* locus. The weakening of H3K27me3 levels in response to knocking down *toy* may be responsible for the elevation of *vg* expression levels. We note that while H3K27me3 levels are strongly reduced in toyKD discs, it is not sufficient, on its own, to significantly alter either *vg* expression or the fate of the eye. It is possible that the reductions in toy expression reduce H3K27me3 levels to a sensitized threshold that can be further modified by the additional loss of PcG factors. How would Toy influence H3K27me3 levels? One exciting and intriguing possibility that requires further experimentation is that Toy could participate in the recruitment of PcG proteins to PRE elements. If this were to be the case it would imply that Toy may provide some specificity to PcG protein binding within the genome of eye-antennal disc. It would also implicate Toy in the direct repression of wing fate.

Lastly, we addressed the question of whether there are multiple paths for the eye to transform into a wing. As we describe in the introduction to this paper, eight different genetic combinations that induce an eye-to-wing transformation are found in the published literature and we present several additional genetic combinations that do the same. It is unclear if all of these genetic combinations have the same transcriptomic effect. We do not address all known genotypes that induce the transformation of the eye into a wing but instead focus on PcKD, toy-PcKD, toy-SfmbtKD, and toy-ScmKD combinations. Our transcriptomic analysis indicates that *vg* expression is ectopically upregulated in all but toy-ScmKD discs. As we have noted above, this suggests that the eye to wing transformation observed in PcKD, toy-PcKD, and toy-SfmbtKD discs are likely occurring through the same molecular mechanism – namely through the ectopic formation of the Vg-Sd complex. In contrast, the eye transforms into a wing in toy-Scm double knockdowns despite the lack of *vg* expression. This suggests that that a second molecular pathway, one that is independent of the Vg-Sd complex, may be engaged in the eye primordium when *toy* and *Scm* are simultaneously knocked down. Our transcriptomic analysis of toy-ScmKD discs indicates that numerous Hox genes as well as the wing selector gene *ap* are differentially expressed. It is a possible that a combination of these factors drives the eye to transform into a wing. The potential existence of multiple molecular paths is consistent with Waddington’s epigenetic landscape model (Waddington, 1957) in which an individual tissue can deviate from its original trajectory and embark on several distinct alternate paths as it undergoes transdetermination and/or transdifferentiation.

## Acknowledgements

We would like to thank Georg Halder (Katholic University, Leuven, Belgium) for the DE-GAL4 strain, the Bloomington Drosophila Stock Center for additional fly strains listed in the materials and methods, the Developmental Studies Hybridoma Bank for antibodies, and the Indiana University Center for Genomics and Bioinformatics for sequence library preparation and next generation sequencing.

## Materials and Methods

### Genetics

The following fly stocks were used from the Bloomington Drosophila Stock Center (BDSC), Indiana University, Bloomington, USA for the main text experiments: UAS-Pho RNAi BL42926, UAS-Calypso RNAi BL56888, UAS-Sce RNAi BL67924, UAS-Sfmbt RNAi BL32473 and BL28677, UAS-Scm RNAi BL55278, UAS-Pc RNAi BL33964, UAS-toy RNAi BL33679, UAS-KASH GFP BL92582, *vg^1^* BL432, and dorsal eye-GAL4 (De-GAL4, Georg Halder, Katholic University, Leuven, Belgium). Flies were grown on standard cornmeal food and all experiments were conducted at 25°C. For scoring of all eye-antennal discs and adult phenotypes, n=30.

### Immunofluorescence and imaging

*Drosophila* eye-antennal imaginal discs were prepared as previously described (Spratford and Kumar, 2014). The following antibodies were used: rat anti-Elav (Developmental Studies Hybridoma Bank [DSHB] #7E8A10, 1:100), mouse anti-Antp (DSHB #8C11, 1:100), FITC-conjugated donkey anti-rat (Jackson #712-095-153, 1:100), Cy3-conjugated donkey anti-mouse (Jackson #715-165-151), Alexa Fluor 647-conjugated phalloidin (Thermo #A22287, 1:20), Rhodamine phalloidin (Thermo #R415, 1:100). Light microscope fluorescent images were taken using a Zeiss Axioplan II compound microscope and processed by Fiji/ImageJ (Schindelin et al., 2012) and Adobe Photoshop software. Adult flies were imaged with a Zeiss Discovery V12 microscope.

### Molecular biology

#### RNA-seq

Samples were collected from eye-antennal at either 120hr or 144hr after egg lay. 50 eye-antennal imaginal discs were dissected from each genotype as previously described (Spratford and Kumar, 2014). Total RNA was extracted with the RNeasy Plus Mini Kit (Qiagen #74134) per kit protocol. 0.3ug RNA was used to generate a polyA-strand specific library with the Illumina TruSeq Stranded mRNA library prep kit. Libraries were prepared and sequenced by the Center for Genomics and Bioinformatics (CGB, Indiana University, Bloomington, USA).

#### CUT&RUN

Chromatin was extracted and purified from ten third-instar eye-antennal or wing imaginal discs as previously described (Weasner et al., 2023). Samples were treated with rabbit anti-trimethyl-histone H3 (Lys27) (C36B11) (Cell Signaling #9733, 1:51) and Normal Rabbit IgG (Cell Signaling #2729, 1:51) antibodies. Cleaved chromatin fragments were purified with the MinElute PCR Purification Kit (Qiagen #28004) per kit protocol. Libraries were prepared and sequenced by the Center for Genomics and Bioinformatics (CGB, Indiana University, Bloomington, USA).

### Computational Analysis

#### RNA-seq

Read quality was assessed with *fastqc* (Andrews, 2010) (v0.11.9). Genome indices were generated via *STAR* (Dobin et al., 2013) and were aligned to the dm6 *Drosophila* genome (with the additional setting *--genomeSAindexNbases* 13 for *Drosophila*). Aligned reads were counted using *Subread* (Liao et al., 2019) function f*eatureCounts* with the settings *-F GTF -t exon -g gene_id --minOverlap 10 -- largestOverlap --primary -s 2 -T*. bigWig files of RNA-seq counts were generated with *deepTools* (Ramirez et al., 2016) (v3.5.1) function *bamCoverage* with the settings *--normalizeUsing CPM -bs 1 -- smoothLength 25 --filterRNAstrand [forward or reverse] -p 16*. Downstream analysis was performed in Rstudio (v4.2.1). Differential expression analysis was performed with *DESeq2* (Love et al., 2014) and heatmaps were generated with *ComplexHeatmap* (Gu et al., 2016). Differentially expressed transcripts were filtered with *dplyr* package (Wickham et al., 2019) to select for enriched/depleted genes in each comparison. Candidate overlap between each time point were then compared with *BioVenn* web application (Hulsen et al., 2008). Gene ontology analysis was performed with *clusterProfiler* (Yu et al., 2012) *eGO* function (settings: *OrgDb = "org.Dm.eg.db", ont = "BP", pAdjustMethod = "fdr", keyType = "SYMBOL"*) or *gprofiler2* (Kolberg et al., 2020). Additional scripts for downstream analysis have been deposited in GitHub (https://github.com/brownhe717/Brown_etal_2024).

#### CUT&RUN

Read quality was assessed with fastqc (Andrews, 2010) (v0.11.9). Genome indices were generated via *Bowtie2* (Langmead et al., 2009) and were aligned to the *Drosophila* dm6 genome and *E. coli* EB1 genomes (with the settings *min_fragment = 10 max_fragment = 700*). The quality and consistency of raw aligned reads were assessed with deeptools (v3.5.1) function *multiBamSummary* and *plotCorrelation* (with the settings *--whatToPlot heatmap --corMethod spearman --plotNumbers*). The number of *E. coli* aligned reads were accessed with *samtools* (Danecek et al., 2021) (v1.15.1) command *view -c -F 4* and used to generate an appropriate scale factor with the calculation: 10,000/[# *E. coli* aligned reads]. Normalized bigWig and bedgraph files of *Drosophila* CUT&RUN reads were generated with *deepTools* (Ramirez et al., 2016) (v3.9/3.5.1) function *bamCoverage* with the corresponding *-- scaleFactor and --outFileFormat [bigwig or bedgraph]* setting. Initial visualization of bigWig reads was carried out in Integrative Genomics Viewer (v2.8.0); figures of chromatin tracks were generated using *Gviz* (Hahne and Ivanek, 2016) in Rstudio (v4.2.1). Additional scripts for downstream analysis have been deposited in GitHub (https://github.com/brownhe717/Brown_etal_2024).

## Data availability

Scripts used for the initial RNA-seq (https://github.com/rpolicastro/RNAseq) and CUT&RUN (https://github.com/gzentner/ChIPseq) pipelines can be found on GitHub (https://github.com/brownhe717/Brown_etal_2024). Raw data files have been deposited in Sequence Read Archive (SRA) under accession number PRJNA1124934.

