## supplemental information for "Differential regulation of eye specification in *Drosophila* by Polycomb Group (PcG) epigenetic repressors"

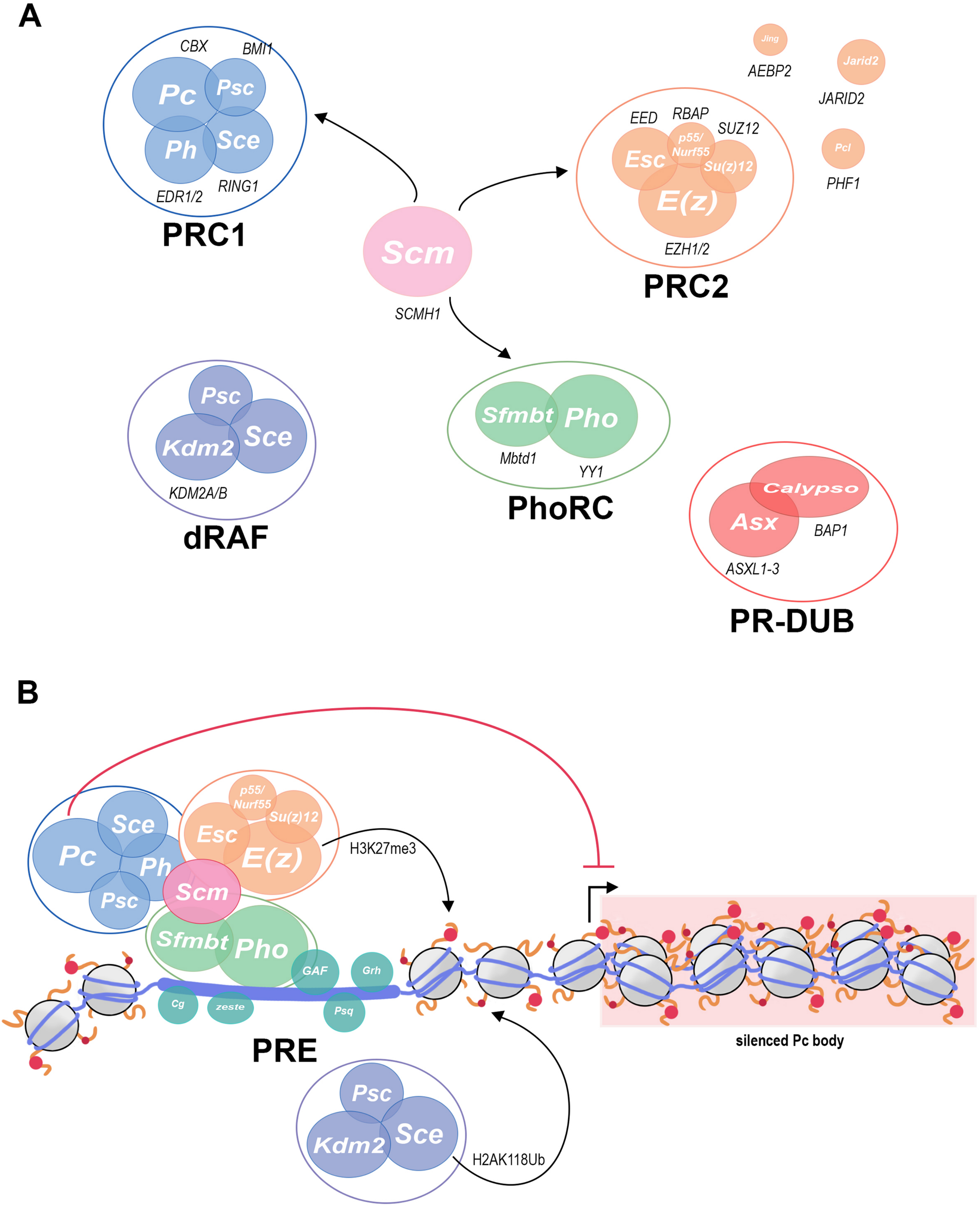
A
CBX
BMI1
Pc
Psc
Ph
Sce
EDR1/2
RING1
PRC1
Scm
SCMH1
Jing
AEBP2
Jarid2
JARID2
EED
RBAP
SUZ12
Esc
p55/ Nurf55
Su(z)12
E(z)
EZH1/2
Pcl
PHF1
PRC2
Psc
Kdm2
Sce
KDM2A/B
dRAF
Sfmbt
Pho
Mbtd1
YY1
PhoRC
Calypso
Asx
BAP1
ASXL1-3
PR-DUB
B
Sce
Pc
Ph
Psc
p55/ Nurf55
Su(z)12
Esc
E(z)
Scm
Sfmbt
Pho
GAF
Grh
Cg
zeste
Psq
H3K27me3
PRE
silenced Pc body
Psc
Sce
Kdm2
H2AK118Ub

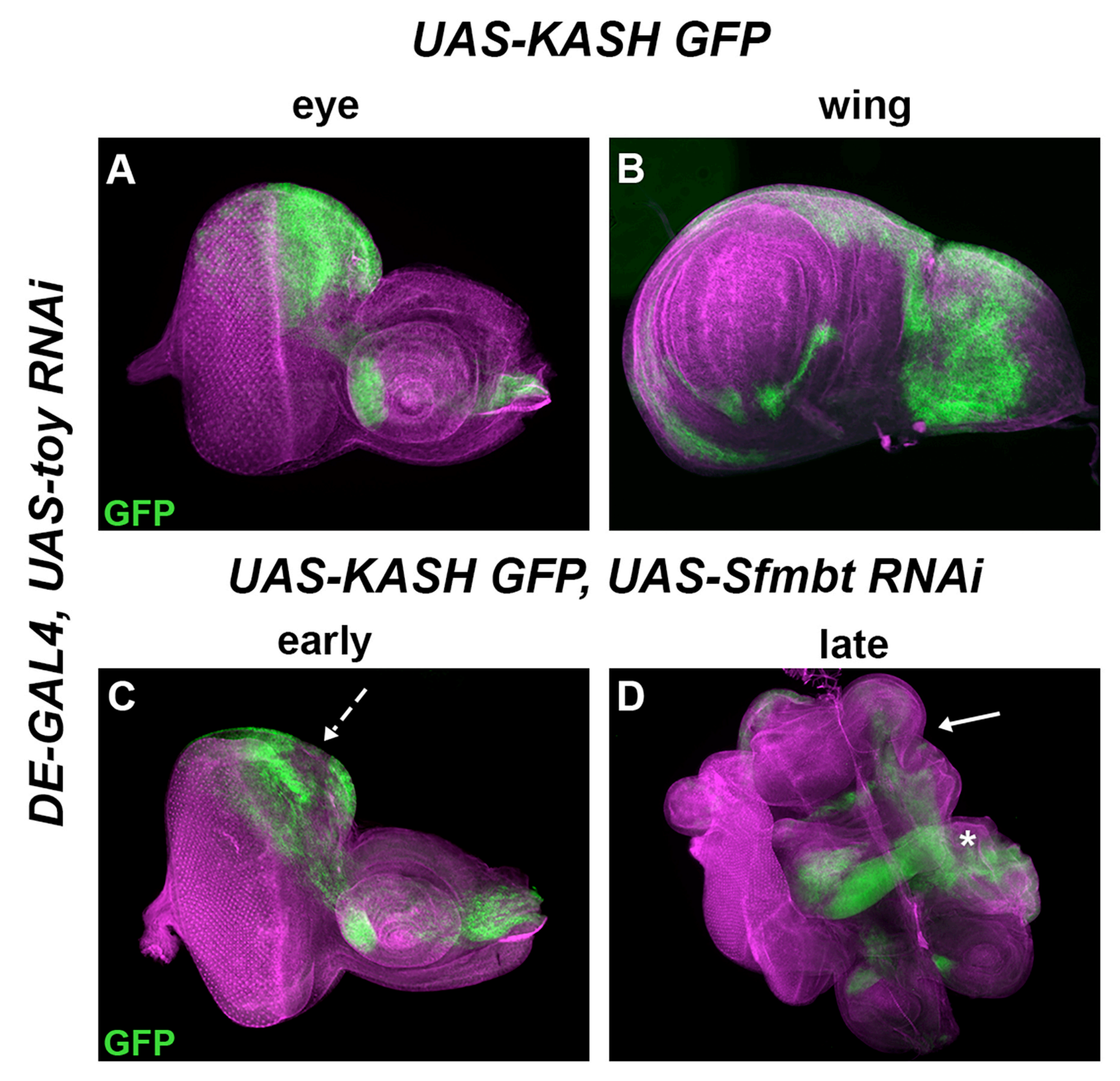

UAS-KASH GFP
eye
wing
DE-GAL4, UAS-toy RNAi
A
GFP
B
UAS-KASH GFP, UAS-Sfmbt RNAi
early
late
C
GFP
D
*

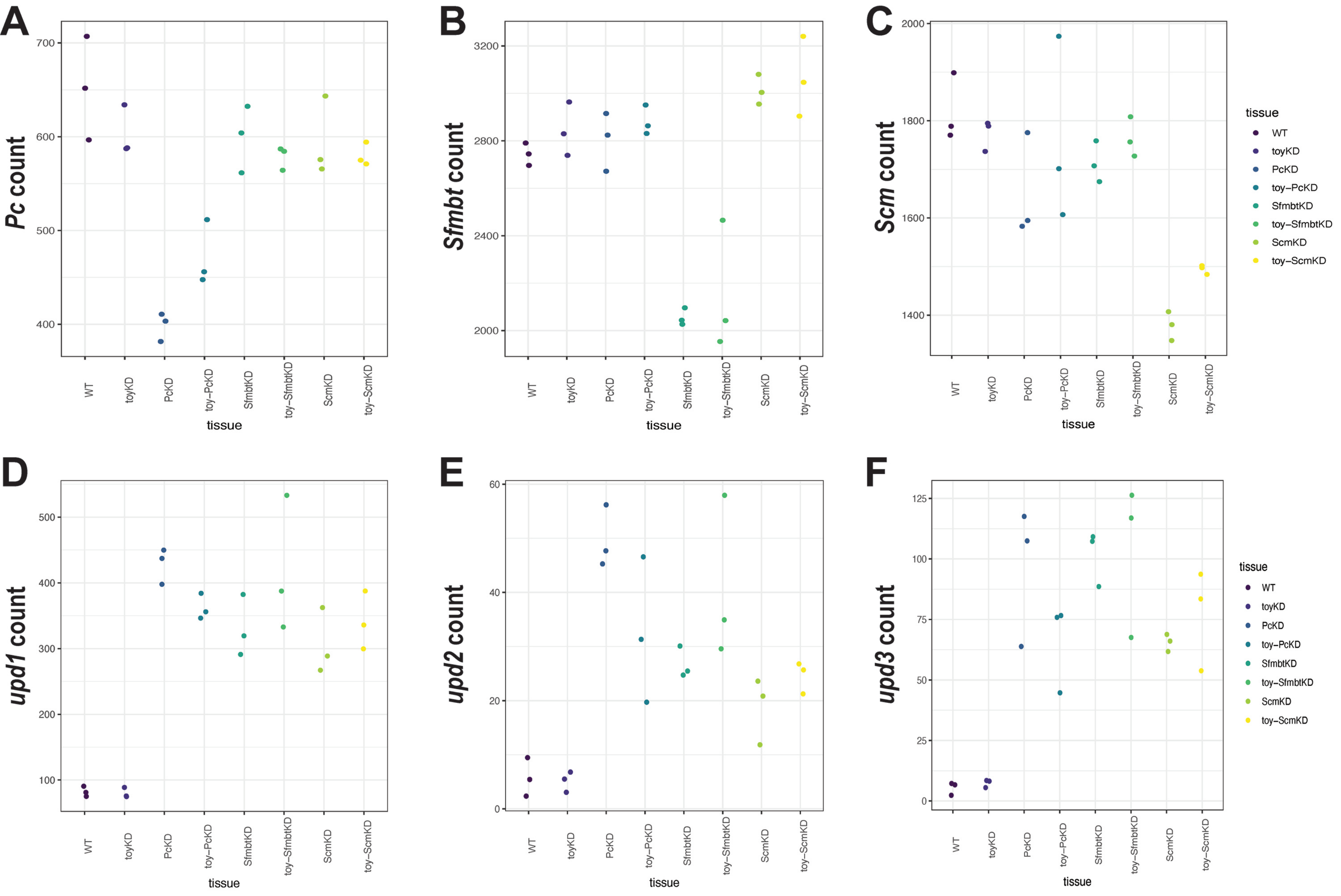
A
B
C
D
E
F
Pc count
Sfmbt count
Scm count
upd1 count
upd2 count
upd3 count
tissue
WT
toyKD
PcKD
toy–PcKD
SfmbtKD
toy–SfmbtKD
ScmKD
toy–ScmKD

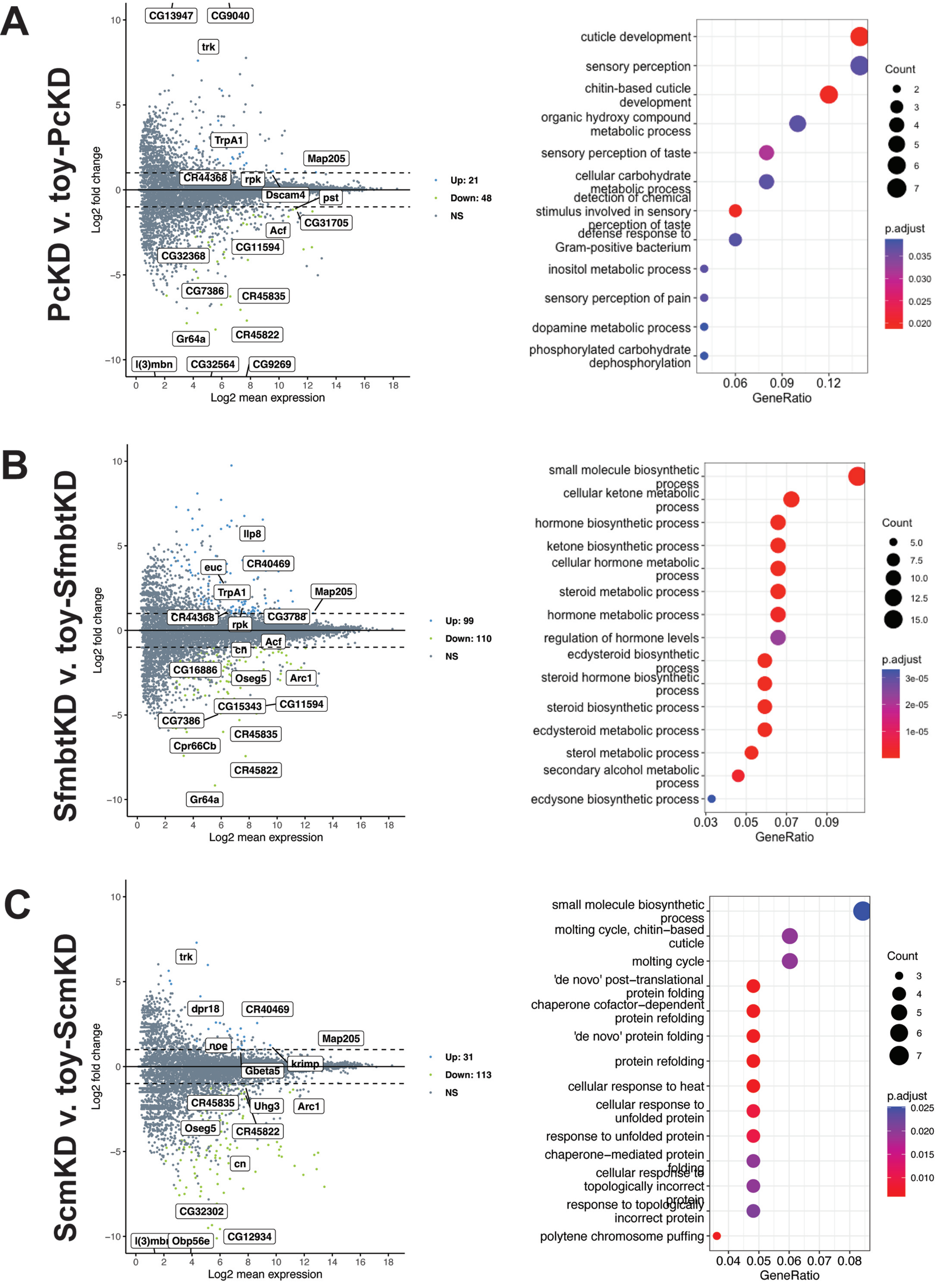
A
PcKD v. toy-PcKD
CG13947
CG9040
trk
TrpA1
Map205
CR44368
rpk
Dscam4
pst
CG31705
Acf
CG11594
CG32368
CG7386
CR45835
Gr64a
CR45822
l(3)mbn
CG32564
CG9269
Log2 fold change
Log2 mean expression
Up: 21
Down: 48
NS
cuticle development
sensory perception
chitin-based cuticle development
organic hydroxy compound metabolic process
sensory perception of taste
cellular carbohydrate metabolic process
detection of chemical stimulus involved in sensory perception of taste
defense response to Gram-positive bacterium
inositol metabolic process
sensory perception of pain
dopamine metabolic process
phosphorylated carbohydrate dephosphorylation
GeneRatio
Count
p.adjust
B
SfmbtKD v. toy-SfmbtKD
Ilp8
euc
CR40469
TrpA1
Map205
CR44368
CG3788
rpk
Acf
cn
CG16886
Oseg5
Arc1
CG15343
CG11594
CG7386
CR45835
Cpr66Cb
CR45822
Gr64a
Up: 99
Down: 110
NS
small molecule biosynthetic process
cellular ketone metabolic process
hormone biosynthetic process
ketone biosynthetic process
cellular hormone metabolic process
steroid metabolic process
hormone metabolic process
regulation of hormone levels
ecdysteroid biosynthetic process
steroid hormone biosynthetic process
steroid biosynthetic process
ecdysteroid metabolic process
sterol metabolic process
secondary alcohol metabolic process
ecdysone biosynthetic process
C
ScmKD v. toy-ScmKD
trk
dpr18
CR40469
Map205
noe
krimp
Gbeta5
CR45835
Uhg3
Arc1
Oseg5
CR45822
cn
CG32302
l(3)mbt
Obp56e
CG12934
Up: 31
Down: 113
NS
small molecule biosynthetic process
molting cycle, chitin–based cuticle
molting cycle
'de novo' post–translational protein folding
chaperone cofactor–dependent protein refolding
'de novo' protein folding
protein refolding
cellular response to heat
cellular response to unfolded protein
response to unfolded protein
chaperone–mediated protein folding
cellular response to topologically incorrect protein
response to topologically incorrect protein
polytene chromosome puffing

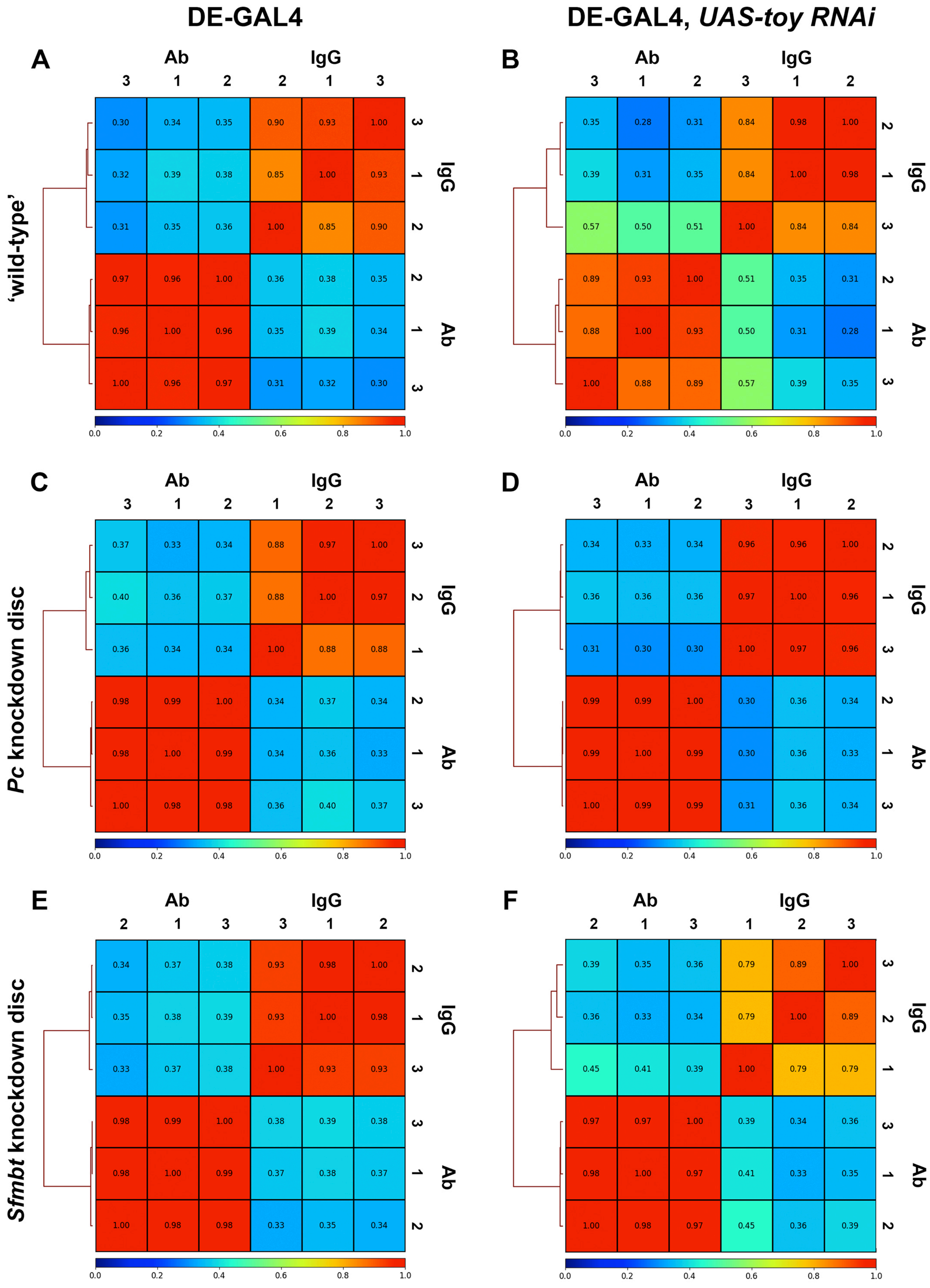
DE-GAL4
DE-GAL4, UAS-toy RNAi
'wild-type'
Pc knockdown disc
Sfmbt knockdown disc
A
Ab IgG
3 1 2 2 1 3
0.30 0.34 0.35 0.90 0.93 1.00
0.32 0.39 0.38 0.85 1.00 0.93
0.31 0.35 0.36 1.00 0.85 0.90
0.97 0.96 1.00 0.36 0.38 0.35
0.96 1.00 0.96 0.35 0.39 0.34
1.00 0.96 0.97 0.31 0.32 0.30
3 1 2 IgG
2 1 3 Ab
0.0 0.2 0.4 0.6 0.8 1.0
B
Ab IgG
3 1 2 3 1 2
0.35 0.28 0.31 0.84 0.98 1.00
0.39 0.31 0.35 0.84 1.00 0.98
0.57 0.50 0.51 1.00 0.84 0.84
0.89 0.93 1.00 0.51 0.35 0.31
0.88 1.00 0.93 0.50 0.31 0.28
1.00 0.88 0.89 0.57 0.39 0.35
2 1 3 IgG
2 1 3 Ab
0.0 0.2 0.4 0.6 0.8 1.0
C
Ab IgG
3 1 2 1 2 3
0.37 0.33 0.34 0.88 0.97 1.00
0.40 0.36 0.37 0.88 1.00 0.97
0.36 0.34 0.34 1.00 0.88 0.88
0.98 0.99 1.00 0.34 0.37 0.34
0.98 1.00 0.99 0.34 0.36 0.33
1.00 0.98 0.98 0.36 0.40 0.37
3 2 1 IgG
2 1 3 Ab
0.0 0.2 0.4 0.6 0.8 1.0
D
Ab IgG
3 1 2 3 1 2
0.34 0.33 0.34 0.96 0.96 1.00
0.36 0.36 0.36 0.97 1.00 0.96
0.31 0.30 0.30 1.00 0.97 0.96
0.99 0.99 1.00 0.30 0.36 0.34
0.99 1.00 0.99 0.30 0.36 0.33
1.00 0.99 0.99 0.31 0.36 0.34
2 1 3 IgG
2 1 3 Ab
0.0 0.2 0.4 0.6 0.8 1.0
E
Ab IgG
2 1 3 3 1 2
0.34 0.37 0.38 0.93 0.98 1.00
0.35 0.38 0.39 0.93 1.00 0.98
0.33 0.37 0.38 1.00 0.93 0.93
0.98 0.99 1.00 0.38 0.39 0.38
0.98 1.00 0.99 0.37 0.38 0.37
1.00 0.98 0.98 0.33 0.35 0.34
2 1 3 IgG
3 1 2 Ab
0.0 0.2 0.4 0.6 0.8 1.0
F
Ab IgG
2 1 3 1 2 3
0.39 0.35 0.36 0.79 0.89 1.00
0.36 0.33 0.34 0.79 1.00 0.89
0.45 0.41 0.39 1.00 0.79 0.79
0.97 0.97 1.00 0.39 0.34 0.36
0.98 1.00 0.97 0.41 0.33 0.35
1.00 0.98 0.97 0.45 0.36 0.39
3 2 1 IgG
3 1 2 Ab
0.0 0.2 0.4 0.6 0.8 1.0

**Supplemental Figure and Table Legends**

**Figure S1. Polycomb Group (PcG) proteins function in several complexes to silence transcription.** (A) Schematic of the PcG members across five complexes: Polycomb Repressive Complex 1 and 2 (PRC1 and PRC2), Pho Repressive Complex (PhoRC), dRING Associated Factor (dRAF), Polycomb Deubiquitinase Complex (PR-DUB), and the Scm linker protein. PRC2-associated proteins, that modify activity of PRC2, (Jing, Jarid2, and Pcl) are shown next to PRC2. Names of mammalian homologs are listed next to each protein. (B) Schematic representing how each complex interacts to repress transcription. Pho first binds to Pc Response Elements (PREs) with the aid of PRE-binding elements (represented here by GAF, Grh, Cg, Psq, Zeste). Sfmbt then interacts with the linker protein, Scm, to recruit PRC1 and PRC2. E(z) will add H3K27me3 (large red dots) to histone tails (orange) which will be 'read' by Pc to ultimately repress transcription. Sce in dRAF complex adds H2AK118Ub (small red dots) in flies – or H2AK119Ub in vertebrates – to aid in PcG recruitment and repression. Figure modified from Kassis et al. 2017.

**Figure S2. Transformed tissue no longer expresses KASH-GFP.** (A-B) Third-instar *DE-toy RNAi* eye-antennal (A) and wing disc (B) expressing KASH-GFP (green) in the dorsal eye and wing. (C-D) Eye-antennal discs knocking down both *toy* and *Sfmbt*, while driving UAS-KASH GFP in the dorsal eye. Early in the transformation (~120hr AEL), GFP begins to be disrupted in the dorsal eye field (C, hatched arrow). By the time the transformation has progressed (~144hr AEL), KASH-GFP is no longer expressed in the prospective pouch (D, arrow) of the transformed tissue and is instead relegated to the transformed hinge tissue (asterisk).

**Figure S3. PcG members are effectively knocked down by RNAi.** (A-F) Plots depicting raw transcript counts of *Pc* (A), *Sfmbt* (B), and *Scm* (C), *upd1* (D), *upd2* (E), and *upd3* (F) across all tested genotypes.

**Figure S4. Toy-specific changes in the double knockdown discs.** (A-C) MA plots (left) and enriched Gene Ontology (GO) analysis (right) of differentially expressed genes between individual Pc (A), Sfmbt (B), or Scm (C) knockdown discs compared to the corresponding double knockdown eye-antennal discs.

**Figure S5. Read similarity with plotCorrelation.** (A-F) Hierarchically clustered heatmap of Spearman's correlation between samples with histone-specific antibody H3K27me3 to the corresponding IgG control for driver alone (A), toyKD (B), PcKD (C), toy-PcKD (D), SfmbtKD (E), and toy-SfmbtKD (F) eye-antennal discs. Numbers below sample identifiers represent the sample number in each triplicate.

**Table S1. PcG knockdown phenotypic results.** Table provides the genotype of each genetic cross and a brief description of the phenotypic result for each RNAi knockdown experiment conducted in this study.

**Table S2. Differential expression of knockdown eye-antennal discs.** Table depicts the raw DESeq2 output of each comparison: WT v. toy or PcG RNAi (sheets 1-4), toyKD v. toy-PcG RNAi (sheets 5-7), PcKD v. PcG RNAi (sheets 8-9), PcKD v. toy-PcG RNAi (sheets 10-12), SfmbtKD v. toy-SfmbtKD (sheet 13), and ScmKD v. toy-ScmKD (sheet 14).

**Table S3. Common differentially expressed transcripts for each genotype.** Table provides a list of all differentially expressed transcripts unique to *Pc*, *Sfmbt*, and *Scm* (sheet 1) or *toy-Pc*, *toy-Sfmbt*, and *toy-Scm* double knockdown discs (sheet 2) as well as those transcripts shared among the three genotypes.
